# Immunotherapy of CT26 murine tumors is characterized by an oligoclonal response against the AH1 tumor rejection antigen

**DOI:** 10.1101/789784

**Authors:** Philipp Probst, Marco Stringhini, Dario Neri

## Abstract

The possibility to cure immunocompetent mice bearing murine CT26 colorectal tumors using cytokine-based therapeutics allows to study the tumor rejection process at a molecular level. Following treatment with L19-mIL12 or F8-mTNF, two antibody fusion proteins which preferentially concentrate a murine cytokine payload at the tumor site, CT26 tumors could be cured in a process that crucially relies on CD8+ T cells. In both settings, the AH1 peptide (derived from the gp70 envelop protein of murine leukemia virus) acted as the main tumor rejection antigen and ~50% of CD8+ T cells in the tumor mass are AH1-specific after therapy. In order to characterize the clonality of the T cell response after successful antibody-cytokine immunotherapy, we isolated CD8+ T cells from tumors and submitted them to T cell receptor (TCR) sequencing. As expected, different TCR sequences were found in different mice, as these molecules originate from a stochastic rearrangement process. CD8+ T cells featuring the ten most abundant TCR sequences represented more than 60% of total CD8+ T cell clones in the tumor mass, but less than 10% in the spleen. Looking at sorted CD8+ T cells from individual animals, AH1-specific TCRs were consistently found among the most abundant sequences. Collectively, these data suggest that the antitumor response driven by two different antibody-cytokine fusions proceeds through an oligoclonal expansion and activation of tumor-infiltrating CD8+ T cells.

## Introduction

Successful immunotherapy is able to induce durable complete responses (CRs) against certain types of tumors, both in animal models of cancer and in patients. For instance, a subset of immunogenic tumors responds to PD-1 blockade and durable CRs have been described in patients with advanced melanoma (1). Similarly, a proportion of patients with metastatic melanoma or renal cell carcinoma can be cured using high-dose interleukin-2 treatment (2,3). In mice, cures of CT26 colorectal cancer has been described in mice receiving immune checkpoint inhibitors (4), antibody-cytokine fusions (5,6) or a combination of these two treatment modalities (7).

CD8+ T cells play a crucial role in the tumor rejection process (8,9). Highly-mutated tumors (e.g., melanomas and non-small cell lung cancers) typically respond better to immune checkpoint inhibitors and it is likely that neo-antigens (i.e., mutated peptides presented on HLA class I molecules) contribute to tumor rejection (10,11). However, growing experimental evidence indicates that also aberrantly-expressed antigens (originating from “non-coding regions” in the human genome) may contribute to tumor surveillance and cancer rejection (12). Recently, Laumont et al. have described that about 90% of targetable tumor specific antigens in two murine cancer cell lines and seven human primary tumors were derived from non-coding regions (12).

AH1 possibly represents the best characterized tumor rejection antigen in mice. It is derived from the gp70 envelope protein of murine leukemia virus (MuLV), which endogenous in the genome of most laboratory mouse strains (13). AH1 was first described by Huang et al. (14) as the major rejection antigen of the CT26 colorectal cancer cell line and it has since been used as a model antigen to investigate CD8^+^ T cell immunity in different mouse tumor cell lines (15).

Our group has previously shown that immunocompetent mice bearing murine tumors can be cured using certain antibody-cytokine fusions with tumor-homing properties. Two antibodies (F8 and L19, specific to the alternatively-spliced EDA and EDB domains of fibronectin, respectively) were found to selectively localize to solid tumors following intravenous administration (16,17) and were used to deliver various types of cytokine payloads to the tumor environment (18). F8-mTNF, a fusion protein featuring murine TNF as pro-inflammatory payload, exhibits an efficient and selective homing to the tumor site after intravenous injection, as revealed by quantitative biodistribution studies with radiolabeled protein preparations (19). Potent therapeutic activity of the immunocytokine has been observed in different immunocompetent mouse models of cancer, when used as a single agent or in combination chemotherapy and peptide vaccines (5,20). Within the neoplastic mass, F8-TNF can induce a rapid necrosis, turning the tumor into a black, hemorrhagic mass. L19-mIL12, bearing murine interleukin-12 as immunostimulatory moiety, was able to cure immunocompetent BALB/c mice with CT26 colorectal cancer or other tumor types (21). In both cases, FACS analysis of tumor-resident CD8+ T cells revealed that AH1-specific T cells represented the majority of lymphocyte specificity within the neoplastic mass, both at the onset of therapy and after a successful therapeutic intervention. Those CD8+ T cells had an exhausted phenotype, being in most cases positive for PD-1, TIM-3 and LAG-3 (5). However, therapeutic results as well as previously published lymphocyte depletion experiments by our group (20) indicate that these AH1-specific CD8+ T cells are not totally inert and may exert lytic functions.

In this article, we aimed at characterizing the features of tumor-infiltrating CD8+ T cells, isolated from neoplastic masses in mice which had been treated with F8-mTNF, L19-mIL12 or with saline. The TCR sequences were, in most cases, different from mouse to mouse, as they originated from the random joining of V, (D) and J segments in T cell precursors. However, we observed an enrichment for certain T cells (as identified by their TCR sequences) and AH1-specific clones featured consistently among the most frequent lymphocytes within the tumor mass. These data suggest that successful immunotherapy of CT26 clones essentially proceeds with an oligoclonal response and that a relatively small number of CD8+ lymphocytes can drive the tumor rejection process.

## Materials and Methods

### Animals and tumor models

CT26 colon carcinoma cells were purchased from the American Type Culture Collection (ATCC). The cell line was handled according to the protocol of the supplier and kept in culture for no longer than 2 months. Authentication including check of post-freeze viability, growth properties and morphology, test for mycoplasma contamination, isoenzyme assay and sterility test were performed by the cell bank before shipment. Eight-week-old female BALB/c mice were obtained from Charles River (Germany). All animal experiments were performed under a project license granted by the Veterinäramt des Kantons Zürich, Switzerland (04/2018)

### Immunocytokine treatment

Exponentially growing CT26 tumor cells were harvested, repeatedly washed and resuspended in saline prior to injection. Cells were implanted subcutaneously (s.c.) in the right flank of the mice using 3×10^6^ cells per animal. Tumor volume was calculated as follows: (length [mm] x width [mm] x width [mm])/2. When tumors were clearly palpable, mice were randomly divided into the different treatment groups (n = 5). F8-TNF and L19-IL12 were injected intravenously (i.v.) in the lateral tail vein. Mice received two injections of either 2 µg F8-TNF or 12 µg of L19-IL12 every 48 h. Saline treated mice were used as control.

### Generation of AH1 tetramers

Bacterial expression of recombinant murine MHC class I H-2L^d^ and of human β2-microglobulin was performed according to established protocols (20). Refolding of the MHC class I molecules with β2-microglobulin and the AH1 peptide was followed by biotinylation with MBP-BirA and size exclusion chromatography (Superdex S75 10/300 GL, GE Healthcare). Assembled monomers were stored in 16% glycerol at −20 ºC. MHC class I tetramers were produced by addition of PE-conjugated streptavidin (Biolegend) to the monomers at a final molecular ratio of 1:4. The AH1 peptide was ordered from Biomatik.

### Sample preparation for flow cytometry

Spleens and tumors were excised 48 h after the second F8-TNF or L19-IL12 injection. Spleens were minced in PBS, treated with RBC Lysis Buffer (Biolegend), passed through a 40 µm cell-strainer (EASYstrainer, greiner bio-one), repeatedly washed and used directly for cell sorting. Tumors were cut into small pieces and digested in RPMI-1640 (Thermo Fisher, with L-glutamine) containing antibiotic-antimycotic solution (Thermo Fisher), 1 mg/mL collagenase II (Thermo Fisher) and 0.1 mg/mL DNase I (Roche) at 37ºC, 5% CO_2_ for 2 h in an orbital shaker at 110 rpm. The cell suspension was then passed through a 40 µm cell-strainer (EASYstrainer, greiner bio-one), repeatedly washed and immediately used for cell sorting.

### Cell sorting

Spleen and tumor cell suspensions were incubated for 20 min on ice with TruStain fcX (anti-mouse CD16/32, Biolegend). Surface staining was performed with PE-coupled AH1 tetramers and fluorochrome-conjugated antibodies against CD8 (53-6.7) and CD4 (GK1.5) which were both purchased from Biolegend. Cells were stained in PBS containing 0.5% bovine serum albumin (BSA) and 2 mM EDTA for 30 min at 4 ºC. 7-AAD was used for staining of dead cells. Cells were sorted on a FACSAria II (BD).

### Bulk TCR Sequencing

100’000 CD8+ T cells from spleen and tumor cell suspensions of a single mouse (n = 3 per treatment group) were sorted into 96-well plates. Cells were immediately processed and genomic DNA (gDNA) was extracted using the QIAamp® DNA Micro kit (Qiagen) according to the manufacturer’s protocol. DNA quality and concentration were assessed using a NanoDrop 2000c spectrophotometer (Thermo Scientific). gDNA was shipped to Adaptive Biotechnologies (Seattle, WA, USA) for TCRB library preparation and high-throughput sequencing using the immunoSEQ Service for mouse TCRB.

### Single-cell TCR Sequencing

Single CD8+ T cells and single AH1+ CD8+ T cells from spleen and tumor cell suspensions of a single mouse were sorted into 96-well plates containing 2 µL 5x iScript reaction mix (iScript Advanced cDNA Synthesis kit, BioRad), 6 µL mQ and 1 µL 1% Triton-X. After sorting, 1 µL of iScript reverse transcriptase was added to each well and RT-PCR was conducted according to the supplier’s protocol. Nested PCR for amplification of TCRα and TCRβ was performed as described by Dash et al. with slight modifications (22). In brief, cDNA synthesis was performed from single cells using the iScript cDNA Synthesis Kit (Bio-Rad) according to the manufacturer’s instructions in the presence of 0.1% Triton X-100 (Sigma-Aldrich). Following RT, two rounds of multiplex nested PCRs were performed with the GoTaq G2 Green Master Mix (Promega) to amplify the CDR3α and CDR3β transcripts from each cell. The PCR products were analyzed on a 1% agarose gel and sequenced using TRAC or TRBC reverse primers for α and β PCR products, respectively. Sequences were matched with the IMGT database (www.imgt.org).

### Data analysis

The Gini coefficient and the Simpson’s index are two values that have been proposed to describe the diversity of TCR sequencing data. We have previously described to calculation of the Gini coefficient (20) for CD8+ T cell sequencing data. The generation of the Simpson’s index has been described elsewhere (23).

Data were analyzed using Prism 7.0 (GraphPad Software, Inc.). Statistical significances were determined with a regular two-way ANOVA test with the Bonferroni post-test. Data represent means ± SEM. P < 0.05 was considered statistically significant. * = p < 0.05, ** = p < 0.01, *** p = < 0.001, **** = p < 0.0001.

## Results

### CD8+ T cells response against CT26 is highly clonal

The immunocompetent CT26 colon carcinoma model has been classified as a “hot” tumor with a rich lymphocyte infiltrate (24). Successful treatment of CT26-bearing BALB/c mice has been described for different immunotherapeutic agents, including the tumor targeting fusion proteins L19-IL12 and F8-TNF (5,21). In both cases, it has been shown that CD8+ T cells recognizing the tumor antigen AH1 represented the majority of lymphocytes within the neoplastic mass after treatment and that these immune cells are crucial for successful tumor rejection.

To analyze changes in the TCR repertoire induced by L19-IL12 and F8-TNF, tumors and spleens of treated mice were excised 48 h after the second injection of the immunocytokine. 100’000 CD8+ T cells from spleens and tumor cell suspensions of a single mouse (n = 3 per treatment group) were sorted, DNA was extracted and used to perform TCRβ deep sequencing. Between 142 – 2531 and 250 – 14173 unique T cell clones were found in tumor and spleen samples, respectively, based on nucleotide CDR3 sequences. T cell clones were than ranked from highest to lowest frequency for each animal and the group average frequency distribution was calculated. This analysis revealed that the 10 most frequent CD8+ T cell clones made up between 60.2% (PBS treated group) – 70.0% (L19-IL12 treated group) of the whole CD8+ TCR repertoire within the tumor (Figure 1A). By contrast, sequences were more evenly distributed within the spleen. A cumulative frequency of only 3.74% (L19-IL12 treated mice) – 8.81% (F8-TNF treated mice) was reached by the 10 most frequent CD8+ T cell clones. TCR diversity was further analyzed using the inverse Simpson’s index and the Gini coefficient. Both values indicate a high clonality of TCR sequences within the CT26 tumor mass and a low clonality of spleen-derived CD8+ T cells, independent of the treatment modality (Figure 1B-D).

**Figure 1.**
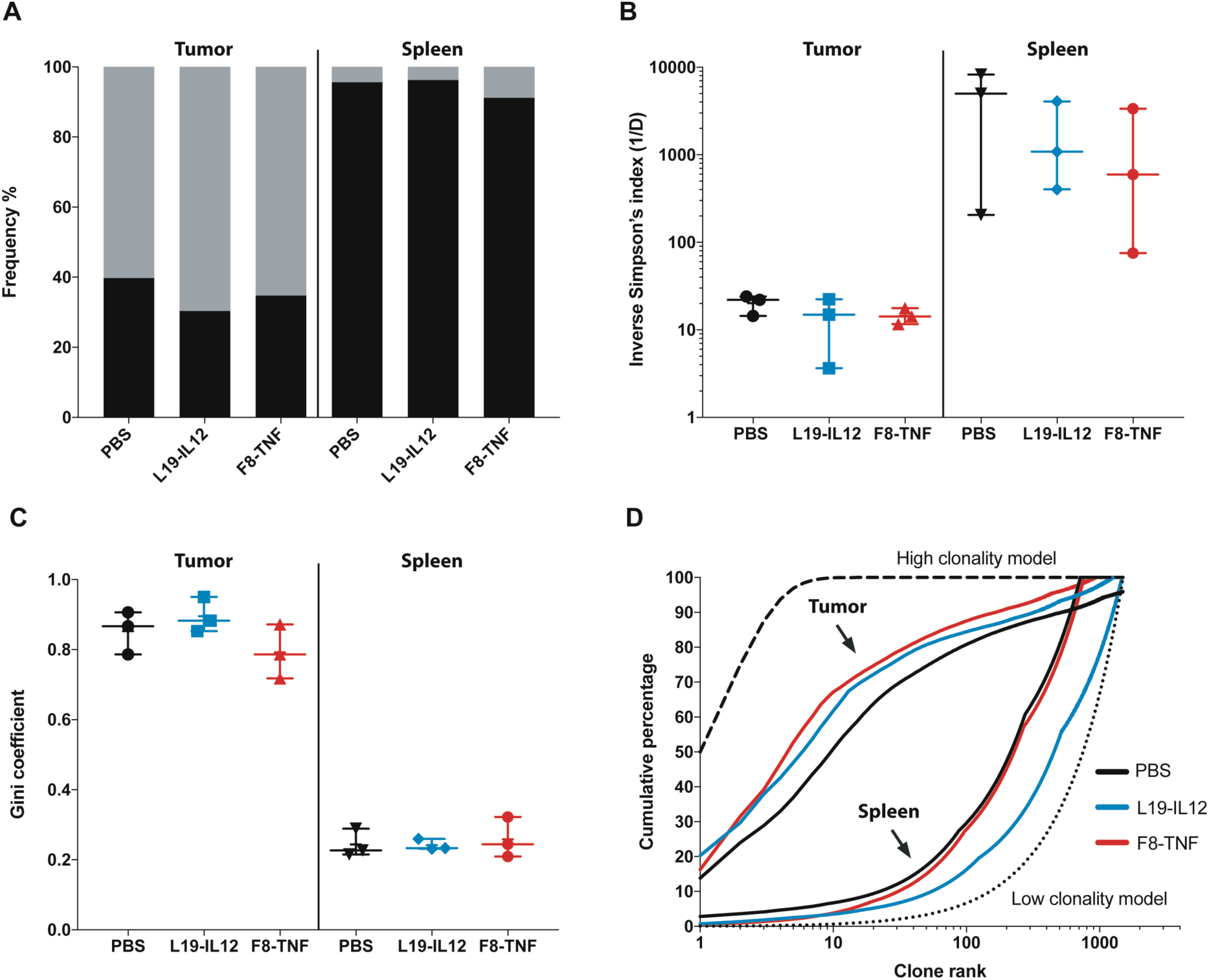
Clonality and frequency distribution of CD8+ T cell clonotypes within tumor and spleen. **A**, Visualization of clone frequency occupancy by the 10 most frequent CD8+ T cell clones based on nucleotide CDR3 sequence. Columns represents the average frequency distribution of the T cell clones within the tumor (left) or within the spleen (right) from animals of the different treatment groups (n = 3). The Top 10 clones are shown in grey, black represents all the other CD8+ T cell clones. **B**, Inverse Simpson’s index values of CD8+ T cells in individuals of the three different treatment groups (n = 3). Treatment groups: PBS (black), L19-IL12 (blue), F8-TNF (red) **C**, Gini index values of CD8+ T cells within the tumor (left) or spleen (right) in mice of the different treatment groups (n = 3). **D**, Average cumulative frequency distribution. CD8+ T cell clones were ranked according to their frequency for each mouse and the average cumulative frequency distribution was calculated for the treatment groups (n = 3). Two models were included in the graph to illustrate a high and a low clonality distribution. For the low clonality model (dotted line), all clones shared a frequency of 0.069% (100%/1450), whereas in the high clonality model (dashed line) a clone with rank n was given a frequency of 50%/2^n-1^.

### Immunocytokine treatment influences the intratumoral TCR repertoire

To gain further insights into the effects of immunocytokine treatment on the TCR repertoire of tumor-infiltrating CD8+ T cells, we determined the frequency of the TRBJ and TRBV segment usage and the CDR3 length distribution for each animal. This analysis revealed a significant increase in the frequency of the TRBJ 01-03 gene segment (Figure 2A) in L19-IL12 treated mice compared to PBS (p = 0.009) and F8-TNF (p = 0.012). In contrast, the usage of TRBJ 02-07 was significantly lower (p = 0.004 compared to PBS and p < 0.0001 compared to F8-TNF treatment). L19-IL12 treated mice also showed an increased usage of the TRBV 02-01 (p = 0.001 compared to F8-TNF group) and 31-01 gene segments (p = 0.035 and p = 0.011 compared to PBS and F8-TNF treatment, respectively) (Figure 2B). A decreased frequency of the TRBV 13-01 gene was observed in immunocytokine treated mice compared to PBS (p = 0.001). Analysis of the CDR3 length distribution revealed no significant change between the three groups with a preferential length of 42 base pairs (Figure 2C).

**Figure 2.**
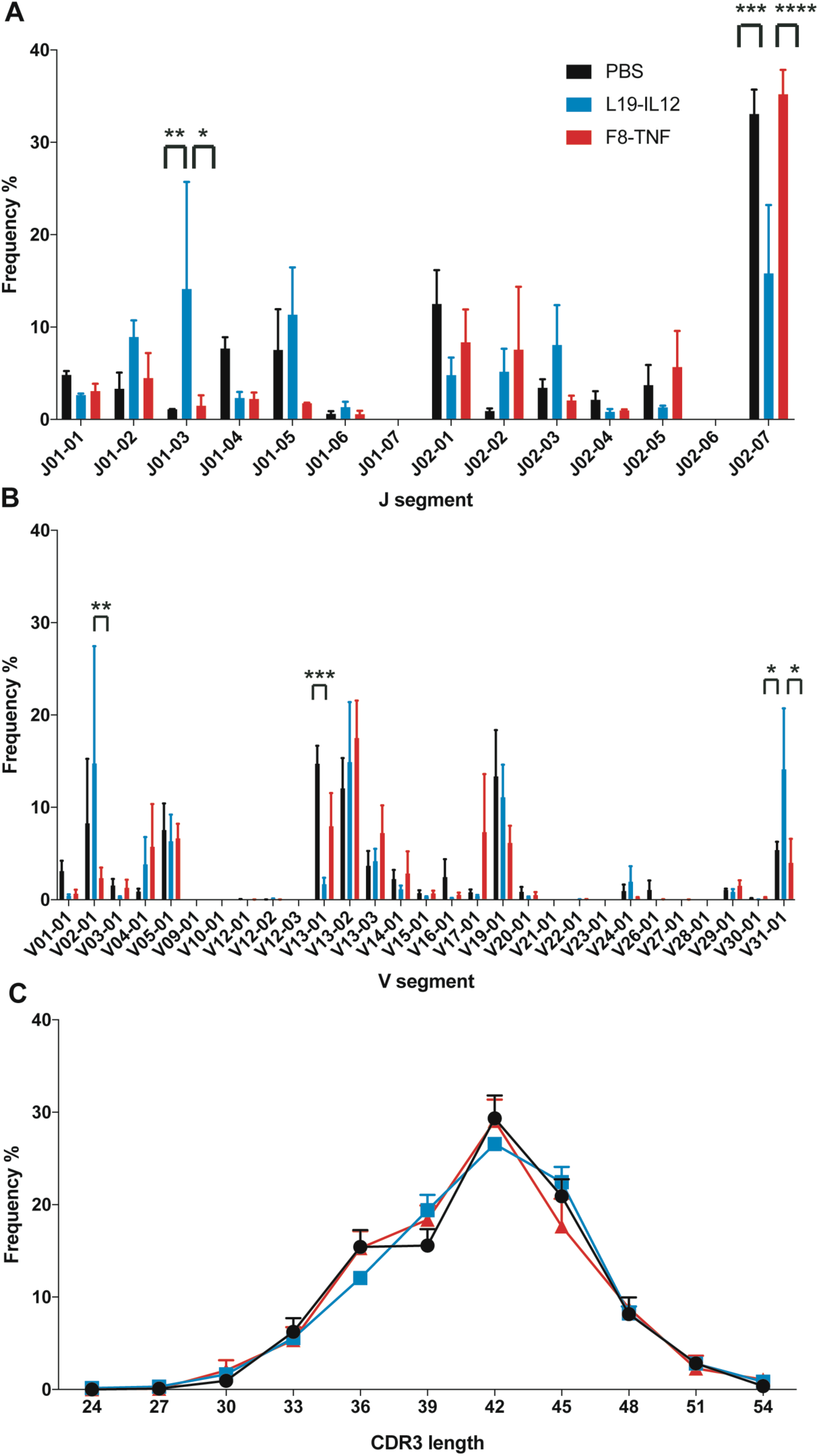
TRBJ and TRBV segment usage, CDR3 sequence length. **A**, Bar plot indicating the average usage of the different TRBJ gene segments in individuals from the different treatment groups (n = 3). Treatment groups: PBS (black), L19-IL12 (blue), F8-TNF (red) **B**, Usage of TRBV gene segments **C**, CDR3 sequence length comparison based on nucleotide sequence. Data represent means ± SEM, n = 3 mice per group. * = p < 0.05, ** = p < 0.01, *** p = < 0.001, **** = p < 0.0001 (regular two-way ANOVA test with the Bonferroni post-test).

### Anti-AH1 response is oligoclonal and driven by highly individual CD8+ T cell clones

To analyze the clonality of the anti-AH1 response, we sorted AH1/H2-L^d^ tetramer–positive CD8+ T cells from spleen and tumor of individual mice and used them for single-cell sequencing of the TCRβ and TCRα region. Sequence analysis revealed that AH1-specific clones featured consistently among the most frequent lymphocytes within the tumor mass of treated (Table 1) and control mice (Table 2). Interestingly, some single clones made up over 20% of the whole TCR repertoire within the tumor (Table 1 and Figure 3A). However, anti-AH1 response was mostly oligoclonal with different clones representing up to 60% of all CD8+ T cells within the tumor. This finding is consistent with previous reports, were a frequency of over 50% of AH1-specific CD8+ T cells was detected in CT26 tumors treated with F8-TNF and a peptide vaccine (5). The majority of CD8+ TCR sequences were unique to one individual and only very few clones were shared between different mice (Figure 3B). We characterized over 100 different AH1 clones using single-cell sequencing and only three clones were shared between more than 2 of the 9 animals. Not a single clone was shared between more than 3 animals. Sequencing results were then used to match AH1-specific TCRβ sequences, which were dominant in an individual animal, with their correspondent TCRα sequence (Table 3).

**Table 1:**
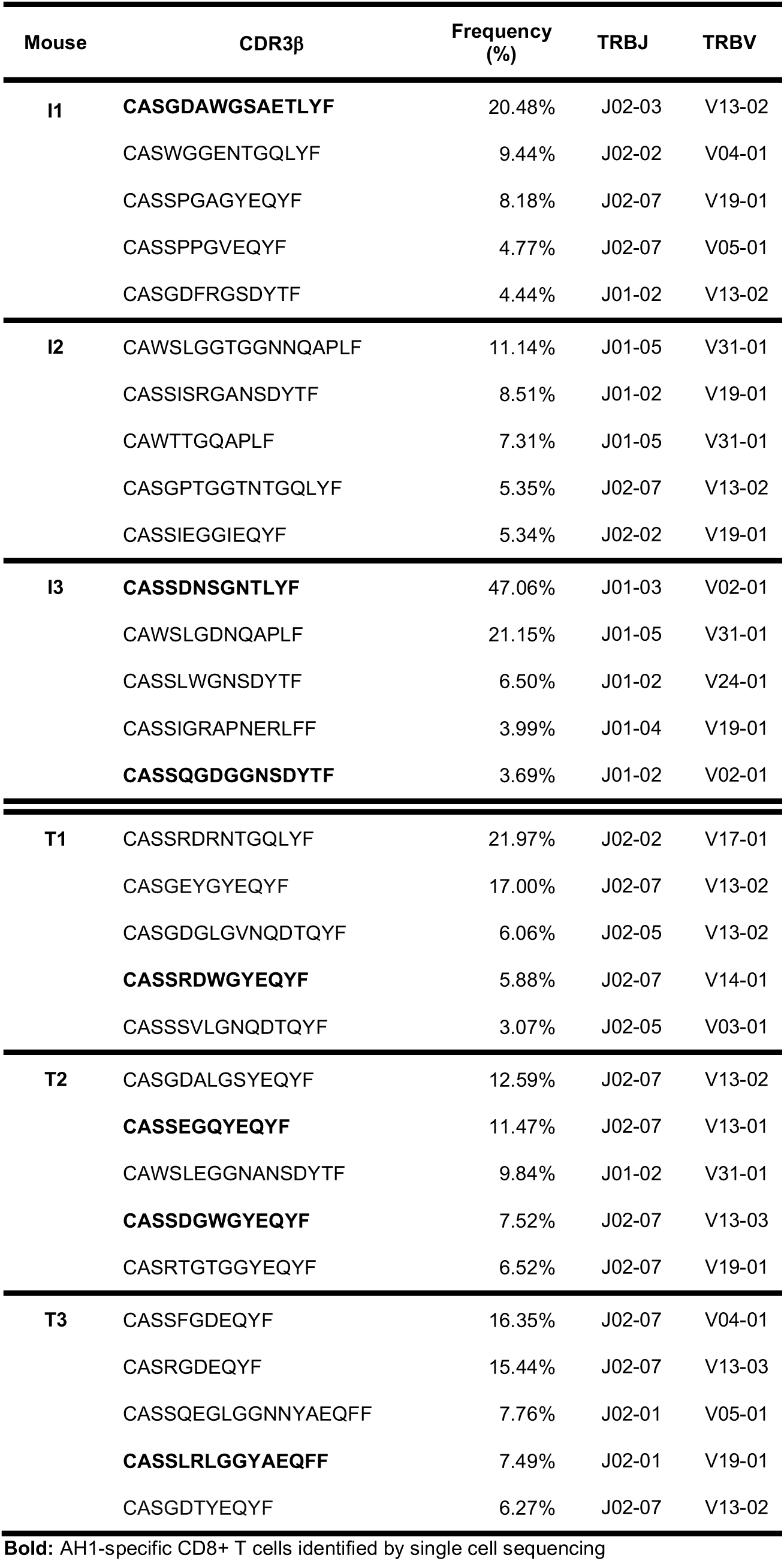
Top five tumor infiltrating T cell clones in each treated animal

**Table 2:** Top five tumor infiltrating T cell clones in each untreated animal

| Mouse | CDR3β | Frequency (%) | TRBJ | TRBV |
| --- | --- | --- | --- | --- |
| **P1** | **CASSSRTGGYAEQFF** | 10.77% | J02-01 | V19-01 |
|  | CASSPRWGGEFEQYF | 8.11% | J02-07 | V05-01 |
|  | CASSDWGGAYEQYF | 7.27% | J02-07 | V16-01 |
|  | CASSERVAQDTQYF | 6.83% | J02-05 | V13-01 |
|  | CTCSAPGHSNERLFF | 5.72% | J01-04 | V01-01 |
| **P2** | CASSPQGAREQYF | 15.82% | J02-07 | V02-01 |
|  | CASGSGTGNQAPLF | 14.20% | J01-05 | V13-02 |
|  | **CASSEGQYEQYF** | 10.72% | J02-07 | V13-01 |
|  | CASSQPGQSNERLFF | 6.71% | J01-04 | V02-01 |
|  | CAWSAGGLYEQYF | 4.36% | J02-07 | V31-01 |
| **P3** | CASSRPGHEQYF | 13.86% | J02-07 | V13-01 |
|  | **CASSIKTGGYAEQFF** | 10.16% | J02-01 | V19-01 |
|  | **CASSLRLGGYAEQFF** | 4.87% | J02-01 | V19-01 |
|  | CASSQDLGLGPSQNTLYF | 4.12% | J02-04 | V05-01 |
|  | CAWSPTGTNNNQAPLF | 3.90% | J01-05 | V31-01 |

**Bold:** AH1-specific CD8<sup>+</sup> T cells identified by single cell sequencing

**Table 3:**
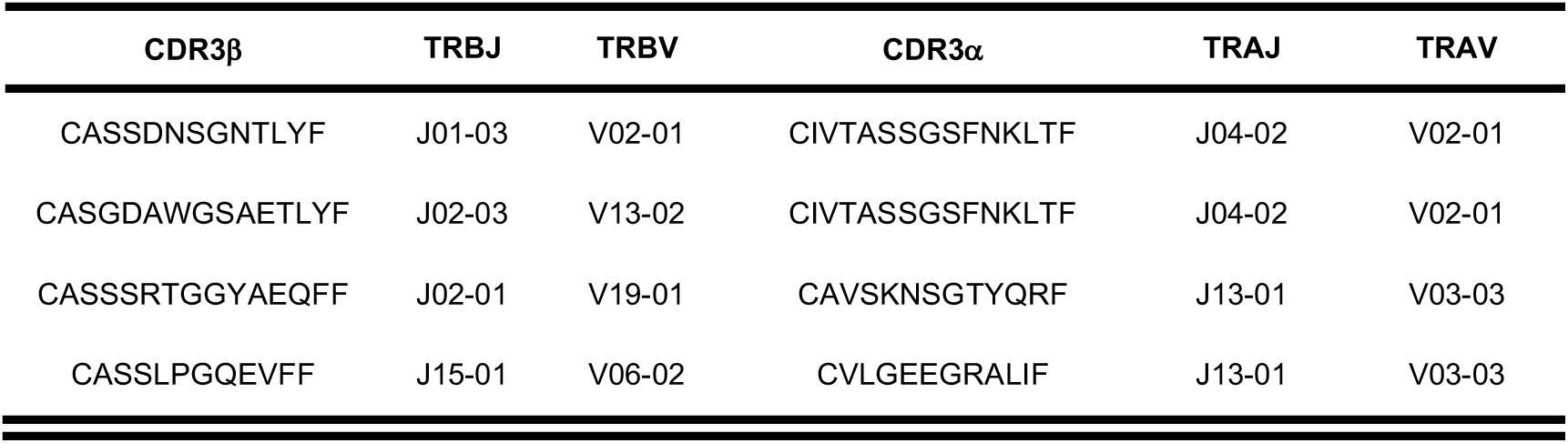
Matching CDR3β and CDR3α sequences of AH1-specific T cells

**Figure 3.**
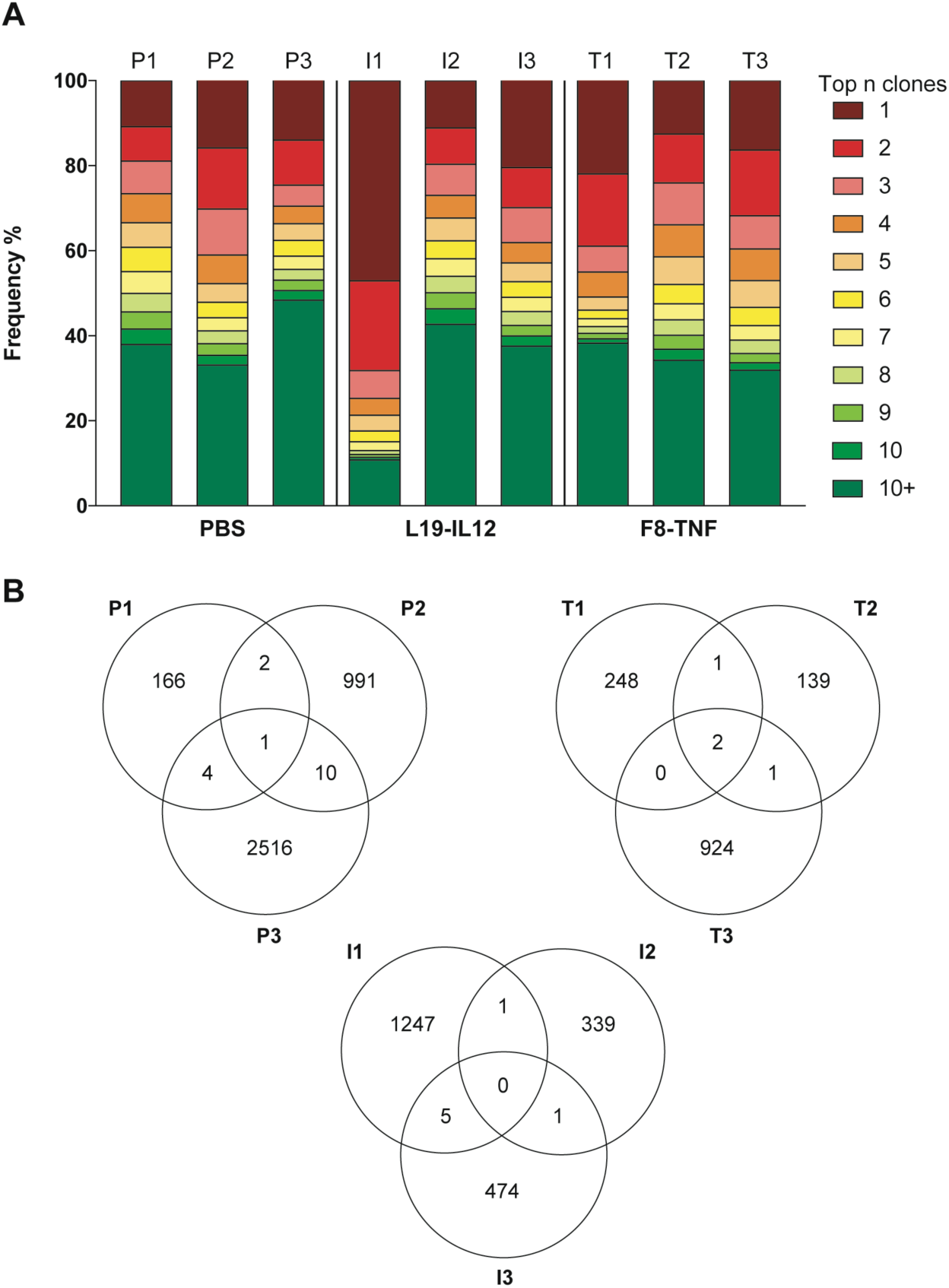
Analysis of the tumor infiltrating CD8+ T cells in each individual animal. **A**, Visualization of the CD8+ T cell clone frequency based on the nucleotide CDR3 sequence. Each column represents the frequency distribution of the T cell clones within the tumor of each individual animal (n = 3 per treatment group). Colors represent the clone ranks. P = PBS, I = L19-IL12, T = F8-TNF. **B**, Venn diagrams of unique tumor infiltrating CD8+ T cells of each individual animal from the respective treatment group (n =3). P = PBS, I = L19-IL12, T = F8-TNF.

## Discussion

We have analyzed CD8+ T cells isolated from tumors and from spleen of mice with CT26 colorectal cancer following therapy with two different antibody-cytokine fusion proteins. TCR sequencing revealed that a relatively small number of T cell clones constitutes the majority of tumor-resident lymphocytes, presumably because of their ability to recognize tumor cells. Indeed, a large proportion of these CD8+ T cells were found to be specific to AH1, a retroviral antigen which is expressed in tumors but not in normal tissues.

Retroviral genes, which have become incorporated into the genome at ancient or more recent times, may represent a substantial proportion of the mammalian genome. For example, it has been estimated that 5% of the murine genome is of retroviral origin (25). Many of these retroviral genes are not annotated as being part of the murine genome and are not expressed in most normal conditions (12,20,26). However, in tumors (and presumably under the influence of elevated interferon-gamma levels) retroviral genes such as the gp70 protein of murine leukemia virus are actively transcribed and give rise to peptides, that can be presented onto MHC class I proteins. Some retroviral antigens are also endowed with transposon activity (27,28).

Rudqvist et al. have recently analyzed the sequence characteristics of T cell clones in BALB/c mice with 4T1 tumors, following radiotherapy treatment and CTLA-4 blockade (29). The authors found that anti-CTLA-4 treatment resulted in fewer T cell clones and more oligoclonal response compared to untreated tumors. Furthermore, the repertoire of T cell receptors reactive with a single tumor antigen was heterogenous but highly clonal, irrespective of the treatment modality (29). In that setting, mice were not cured by the combination treatment, whereas CT26 tumors can be eradicated both by F8-mTNF and by L19-mIL12 (5,30).

In a recent seminal paper, Laumont et al. have reported that noncoding regions may represent the main source of targetable tumor-specific antigens (12). The authors used a combination of RNAseq data (to define actively transcribed genes, including the aberrantly expressed ones) and mass spectrometry-based immunopeptidome analysis to identify putative tumor rejection antigens, which were subsequently functionally validated by vaccination strategies (12). The authors reported that, in addition to AH1, also other peptides were efficiently presented on MHC class I in BALB/c mice and could serve as rejection antigens for CT26. We have recently confirmed that some of those antigens were indeed presented (LPQELPGLVVL, MPHSLLPLVTF and SPHQVFNL) by FACS analysis of T cells with cognate tetramer reagents (Stringhini, Probst and Neri, unpublished observations). However, the frequency of specific T cells was lower, compared to AH1.

We have recently shown that BALB/c mice, which are cured from tumors using certain cytokine-based therapeutics, are able to reject heterologous tumors derived from the same mouse strain (20). The rejection process, which crucially depended on CD8+ T cells, appeared to target the AH1 antigen, since tumor variants that do not present AH1 (e.g., F1F tumors) were not rejected (20).

The observation of an oligoclonal T cell response to CT26 tumors in mice bears some analogy with the T cell response in patients with Merkel Cell Carcinoma (MCC). Miller et al. have reported that polyomavirus-specific T cells may represent up to 18% of the T cell specificities within the tumor mass. Interestingly, an increased number of antigen specific clonotypes correlated with better prognosis (31). Intratumoral infiltration by Merkel Cell polyomavirus-specific T cells was associated with significantly improved MCC-specific survival, suggesting that augmenting the number or avidity of virus-specific T cells may have therapeutic benefit.

Aberrantly-expressed antigens have been reported for human tumors and may be relevant for tumor surveillance. Besides virally-driven tumors (e.g., MCC and HPV-derived malignancies), twenty potentially immunogenic endogenous retroviruses have recently been reported in clear-cell renal cell carcinoma (ccRCC) patients (32). Panda et al. reported that tumors from patients with a high expression of retroviral sequences showed increased immune infiltration and generally responded better to treatment with immune checkpoint inhibitors (32). These findings may explain while ccRCC are immunogenic tumors that respond well to PD-1 blockade (33) and to interleukin-2 (34), in spite of the fact that those tumors exhibit a not particularly high mutation rate (35).

The relative relevance of neo-epitopes (resulting from the MHC class I presentation of mutated peptides) and of aberrantly-expressed antigens for tumor rejection is still a matter of debate. In principle, it would be interesting to perform TCR sequence analysis for mouse models of cancer, which respond to immunotherapy in a process where mutated peptides represent the major source of rejection antigens. Kreiter et al. have reported that CT26 immunotherapy can efficiently be performed using vaccination strategies based on mutated MHC Class II epitopes (36). Similarly, Gubin et al. have shown tumor rejection in mice with mutated MHC Class I antigen (37).

AH1 represents a validated tumor rejection antigen in BALB/c mice and the activation of a relatively small set of T cell clones can eradicate CT26 tumors. It would be interesting to learn whether, in principle a single CD8+ T cell specificity may be sufficient for rejecting tumors in mice, after a suitable expansion and activation. Experiments with AH1-specific clones are currently on-going in our laboratory (M. Stringhini, unpublished results). Cell-based therapeutic strategies may have a clinical relevance. For example, Rosenberg and colleagues have been able to achieve durable complete responses in patients with melanoma and breast cancer, using a judicious selection of tumor-reactive T cells, which were expanded *ex vivo* and re-administered to cancer patients (38,39). The practical implementation of these strategies could greatly benefit from a detailed knowledge of tumor rejection antigens (thus enabling, for example, sorting procedures) and of the minimal clonality of a T cell response which is required for a cancer cure.

The TCR sequence analysis presented in this paper reflects the T cell response in a mouse model of cancer, that can be cured by the targeted delivery of pro-inflammatory cytokines. The CT26 model has been classified as a “hot” tumor, characterized by a rich lymphocyte infiltrate (24). A sequence analysis similar to the one described in this article could be used, in the future, to study the performance of other types of immunotherapy. Moreover, it should be possible to use TCR sequences described in this work for the engineering of stably-transduced T cell clones (40), thus allowing to study this form of cell-based therapy in a molecularly-defined immunocompetent mouse setting. Engineered T cells bearing TCRs as tumor recognition moieties may be more suitable than CAR-T’s for the treatment of solid malignancies, since it is unlikely that antibodies may recognize antigens which are truly tumor-specific.

## Acknowledgments

D. N. gratefully acknowledges ETH Zürich, the Swiss National Science Foundation (Grant Nr. 310030B_163479/1) and the European Research Council (ERC, under the European Union’s Horizon 2020 research and innovation program, grant agreement 670603) for financial support.

## Notes

Conflict of interest disclosure: D.N. is co-founder and shareholder of Philogen, a biotech company that owns the F8 antibody. The authors have no additional financial interests.

## References

1. Schachter J, Ribas A, Long GV, Arance A, Grob J-J, Mortier L, et al. Pembrolizumab versus ipilimumab for advanced melanoma: final overall survival results of a multicentre, randomised, open-label phase 3 study (KEYNOTE-006). The Lancet 2017;390:1853–62

2. Fyfe G, Fisher RI, Rosenberg SA, Sznol M, Parkinson DR, Louie AC. Results of treatment of 255 patients with metastatic renal cell carcinoma who received high-dose recombinant interleukin-2 therapy. 1995;13:688–96

3. Atkins MB, Lotze MT, Dutcher JP, Fisher RI, Weiss G, Margolin K, et al. High-Dose Recombinant Interleukin 2 Therapy for Patients With Metastatic Melanoma: Analysis of 270 Patients Treated Between 1985 and 1993. 1999;17:2105-

4. Hutmacher C, Gonzalo Núñez N, Liuzzi AR, Becher B, Neri D. Targeted Delivery of IL2 to the Tumor Stroma Potentiates the Action of Immune Checkpoint Inhibitors by Preferential Activation of NK and CD8+ T Cells. 2019

5. Probst P, Stringhini M, Ritz D, Fugmann T, Neri D. Antibody-based Delivery of TNF to the Tumor Neovasculature Potentiates the Therapeutic Activity of a Peptide Anticancer Vaccine. Clinical Cancer Research 2019;25:698–709

6. Hemmerle T, Neri D. The antibody-based targeted delivery of interleukin-4 and 12 to the tumor neovasculature eradicates tumors in three mouse models of cancer. Int J Cancer 2014;134:467–77

7. Hutmacher C, Gonzalo Núñez N, Liuzzi AR, Becher B, Neri D. Targeted Delivery of IL2 to the Tumor Stroma Potentiates the Action of Immune Checkpoint Inhibitors by Preferential Activation of NK and CD8^+^ T Cells. 2019

8. Boon T, Cerottini J-C, Van den Eynde B, Van der Bruggen P, Van Pel A. Tumor Antigens Recognized by T Lymphocytes. 1994;12:337–65

9. Reading JL, Gálvez-Cancino F, Swanton C, Lladser A, Peggs KS, Quezada SA. The function and dysfunction of memory CD8+ T cells in tumor immunity. 2018;283:194–212

10. Schumacher TN, Schreiber RD. Neoantigens in cancer immunotherapy. Science 2015;348:69–74

11. Van Allen EM, Miao D, Schilling B, Shukla SA, Blank C, Zimmer L, et al. Genomic correlates of response to CTLA-4 blockade in metastatic melanoma. Science 2015;350:207–11

12. Laumont CM, Vincent K, Hesnard L, Audemard É, Bonneil É, Laverdure J-P, et al. Noncoding regions are the main source of targetable tumor-specific antigens. 2018;10:eaau5516

13. Jenkins NA, Copeland NG, Taylor BA, Lee BK. Organization, distribution, and stability of endogenous ecotropic murine leukemia virus DNA sequences in chromosomes of Mus musculus. Journal of Virology 1982;43:26–36

14. Huang AY, Gulden PH, Woods AS, Thomas MC, Tong CD, Wang W, et al. The immunodominant major histocompatibility complex class I-restricted antigen of a murine colon tumor derives from an endogenous retroviral gene product. Proceedings of the National Academy of Sciences, USA 1996;93:9730–5

15. Miller CT, Graham LJ, Bear HD. Adoptive immunotherapy (AIT) of established tumors with tumor antigen peptide-sensitized T cells. Cancer Research 2004;64:1264-

16. Villa A, Trachsel E, Kaspar M, Schliemann C, Sommavilla R, Rybak JN, et al. A high-affinity human monoclonal antibody specific to the alternatively spliced EDA domain of fibronectin efficiently targets tumor neo-vasculature in vivo. Int J Cancer 2008;122:2405–13

17. Tarli L, Balza E, Viti F, Borsi L, Castellani P, Berndorff D, et al. A high-affinity human antibody that targets tumoral blood vessels. Blood 1999;94:192–8

18. Pasche N, Neri D. Immunocytokines: a novel class of potent armed antibodies. Drug Discov Today 2012;17:583–90

19. Hemmerle T, Probst P, Giovannoni L, Green AJ, Meyer T, Neri D. The antibody-based targeted delivery of TNF in combination with doxorubicin eradicates sarcomas in mice and confers protective immunity. Br J Cancer 2013;109:1206–13

20. Probst P, Kopp J, Oxenius A, Colombo MP, Ritz D, Fugmann T, et al. Sarcoma Eradication by Doxorubicin and Targeted TNF Relies upon CD8+ T-cell Recognition of a Retroviral Antigen. Cancer Research 2017;77:3644–54

21. Puca E, Probst P, Stringhini M, Murer P, Pellegrini G, Cazzamalli S, et al. The antibody-based delivery of interleukin-12 to solid tumors boosts NK and CD8+ T cell activity and synergizes with immune check-point inhibitors. 2019:684100

22. Dash P, McClaren JL, Oguin TH, III, Rothwell W, Todd B, Morris MY, et al. Paired analysis of TCRα and TCRβ chains at the single-cell level in mice. The Journal of Clinical Investigation 2011;121:288–95

23. Kitaura K, Shini T, Matsutani T, Suzuki R. A new high-throughput sequencing method for determining diversity and similarity of T cell receptor (TCR) α and β repertoires and identifying potential new invariant TCR α chains. BMC Immunology 2016;17:38

24. Mosely SIS, Prime JE, Sainson RCA, Koopmann J-O, Wang DYQ, Greenawalt DM, et al. Rational Selection of Syngeneic Preclinical Tumor Models for Immunotherapeutic Drug Discovery. 2017;5:29–41

25. Kassiotis G. Endogenous Retroviruses and the Development of Cancer. The Journal of Immunology 2014;192:1343–9

26. Downey RF, Sullivan FJ, Wang-Johanning F, Ambs S, Giles FJ, Glynn SA. Human endogenous retrovirus K and cancer: Innocent bystander or tumorigenic accomplice? International Journal of Cancer 2015;137:1249–57

27. Chuong EB, Elde NC, Feschotte C. Regulatory evolution of innate immunity through co-option of endogenous retroviruses. Science 2016;351:1083–7

28. Frank JA, Feschotte C. Co-option of endogenous viral sequences for host cell function. Current Opinion in Virology 2017;25:81–9

29. Rudqvist N-P, Pilones KA, Lhuillier C, Wennerberg E, Sidhom J-W, Emerson RO, et al. Radiotherapy and CTLA-4 Blockade Shape the TCR Repertoire of Tumor-Infiltrating T Cells. 2018;6:139–50

30. Puca E, Probst P, Stringhini M, Murer P, Pellegrini G, Cazzamalli S, et al. The antibody-based delivery of interleukin-12 to solid tumors boosts NK and CD8^+^ T cell activity and synergizes with immune check-point inhibitors. 2019:684100

31. Miller NJ, Church CD, Dong L, Crispin D, Fitzgibbon MP, Lachance K, et al. Tumor-Infiltrating Merkel Cell Polyomavirus-Specific T Cells Are Diverse and Associated with Improved Patient Survival. 2017;5:137–47

32. Panda A, de Cubas AA, Stein M, Riedlinger G, Kra J, Mayer T, et al. Endogenous retrovirus expression is associated with response to immune checkpoint blockade in clear cell renal cell carcinoma. JCI Insight 2018;3

33. Rini BI, Plimack ER, Stus V, Gafanov R, Hawkins R, Nosov D, et al. Pembrolizumab plus Axitinib versus Sunitinib for Advanced Renal-Cell Carcinoma. 2019;380:1116–27

34. Rosenberg SA. IL-2: The First Effective Immunotherapy for Human Cancer. 2014;192:5451–8

35. Alexandrov LB, Nik-Zainal S, Wedge DC, Aparicio SA, Behjati S, Biankin AV, et al. Signatures of mutational processes in human cancer. Nature 2013;500:415–21

36. Kreiter S, Vormehr M, van de Roemer N, Diken M, Löwer M, Diekmann J, et al. Mutant MHC class II epitopes drive therapeutic immune responses to cancer. Nature 2015;520:692–6

37. Gubin MM, Zhang X, Schuster H, Caron E, Ward JP, Noguchi T, et al. Checkpoint blockade cancer immunotherapy targets tumour-specific mutant antigens. Nature 2014;515:577

38. Rosenberg SA, Dudley ME. Adoptive cell therapy for the treatment of patients with metastatic melanoma. Current Opinion in Immunology 2009;21:233–40

39. Zacharakis N, Chinnasamy H, Black M, Xu H, Lu Y-C, Zheng Z, et al. Immune recognition of somatic mutations leading to complete durable regression in metastatic breast cancer. Nature Medicine 2018;24:724–30

40. Schumacher TNM. T-cell-receptor gene therapy. Nature Reviews Immunology 2002;2:512–9

